## Supplemental materials for "Impact of anionic lipids on the energy landscape of conformational transition in anion exchanger 1 (AE1)"

### Supporting Information

#### Supplementary Figures

Supplementary Movie 1: **Conformational changes, $HCO_3^-$ transport, and $PIP_2$ interactions during the OF-IF transition in AE1.** In this movie, an extracellular $HCO_3^-$ initially diffuses and binds to R730 of an AE1 protomer in the OF state. The protein then undergoes the OF to IF transition under the applied biases. Concurrently, the salt bridge between $PIP_2$ and K743 is disrupted. Finally, $HCO_3^-$ dissociates from the binding site and is translocated to the cytoplasmic side.

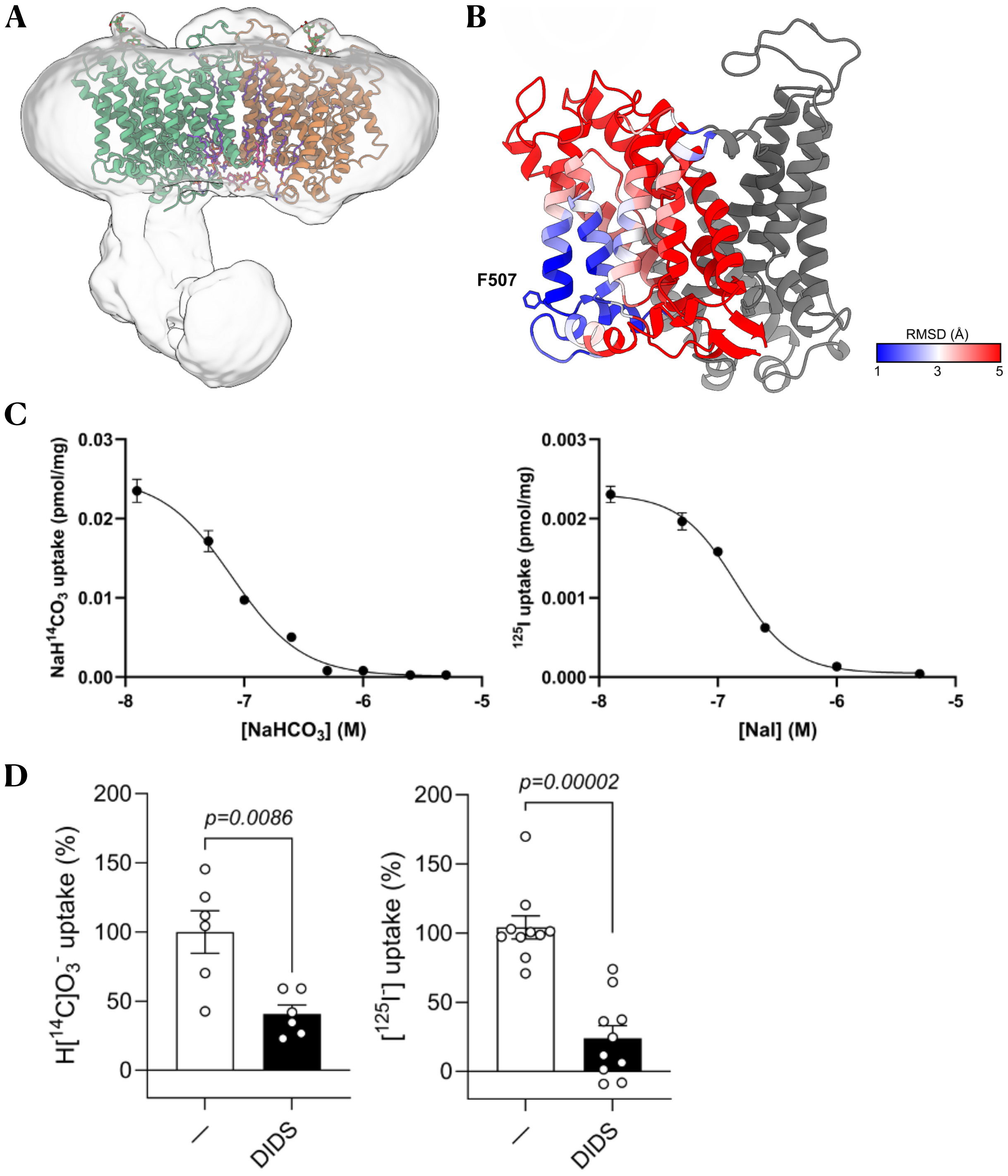


Figure S1: **cryo-EM density and substrate uptake by AE1** (A) AE1 IF1-OF model (IF1 in orange and OF in green) superimposed with the low-threshold map highlights the low-resolution density of the cytosolic domain. (B) Structural alignment between AE1 protomers in IF1 and OF state (colored by RMSD), shows that the movement of the TD doesn't affect the residue F507. (C) AE1-specific 10-s uptake of 10 μM NaH[$^{14}$C]$O_3$ or Na[$^{125}$I] was measured with an isotopic dilution of $NaHCO_3$ or NaI, yielding a log($EC_{50}$) = −7.10 ± 0.02 for $NaHCO_3$ (corresponding to a mean of 79 nM) and log($EC_{50}$) = −6.84 ± 0.01 for NaI (corresponding to a mean of 143 nM). (D) 1-min uptake of 10 μM NaH[$^{14}$C]$O_3$ or Na[$^{125}$I] into proteoliposomes containing AE1 was measured in the presence or absence of 1 mM DIDS.

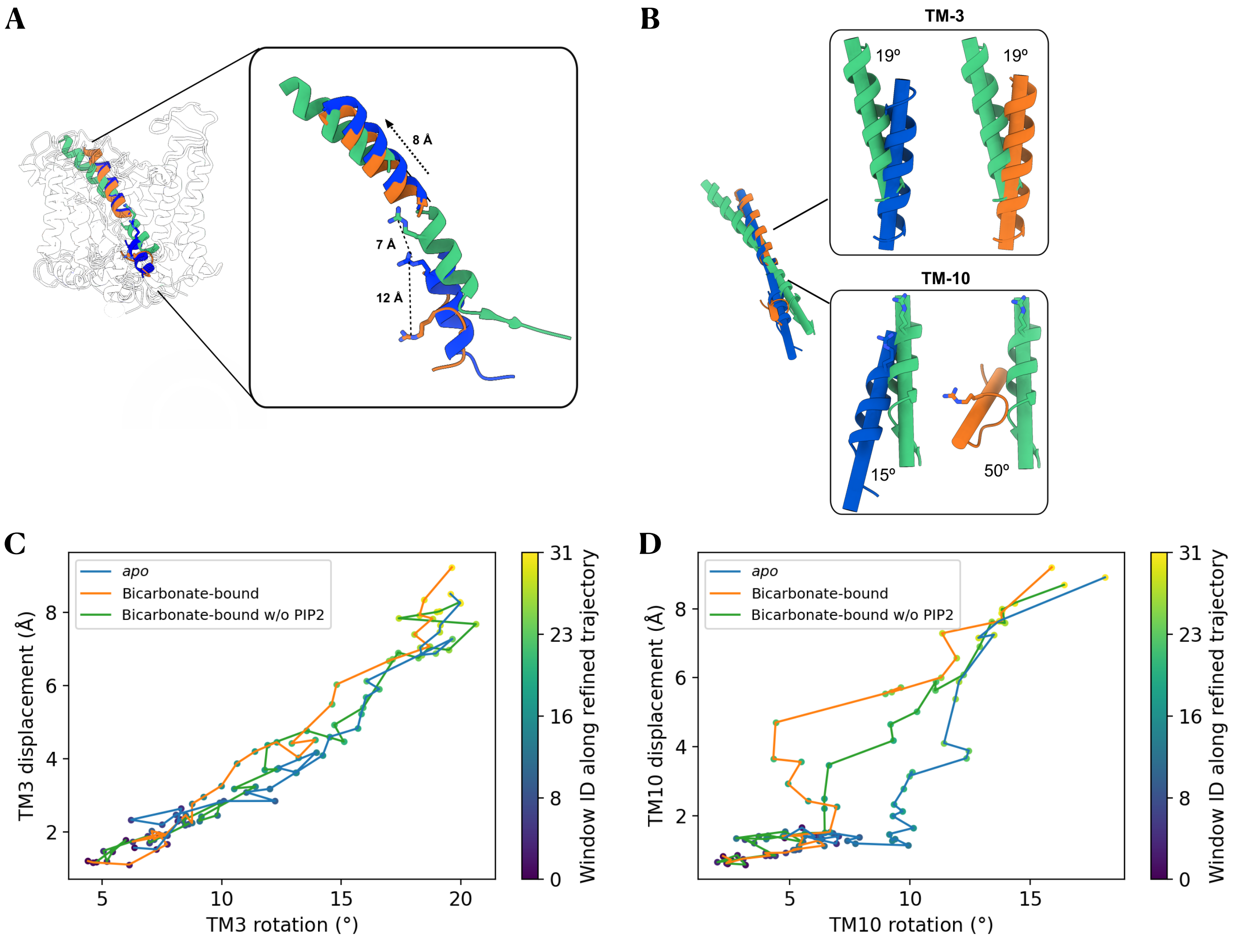


Figure S2: **TM3 and TM10 rotation and displacement.** (A) TM3 shifts by 8 Å between the IF (blue and orange) and OF (green) states, while R730 moves 7 Å between the OF and IF2 (blue) states and 12 Å between the IF2 and IF1 (orange) states. (B) Comparison of the rotation angles between TM3 and TM10 among the three conformational states, with colors matching those shown in A. (C) Calculated rotation and displacement of TM3 along SMwST trajectories for the three simulation systems: *apo*, $HCO_3^-$-bound, and $HCO_3^-$-bound without $PIP_2$. Points represent the averaged center for each window with color from dark blue to yellow indicating windows 0 to 31. (D) Calculated rotation and displacement of TM10 along SMwST trajectories.

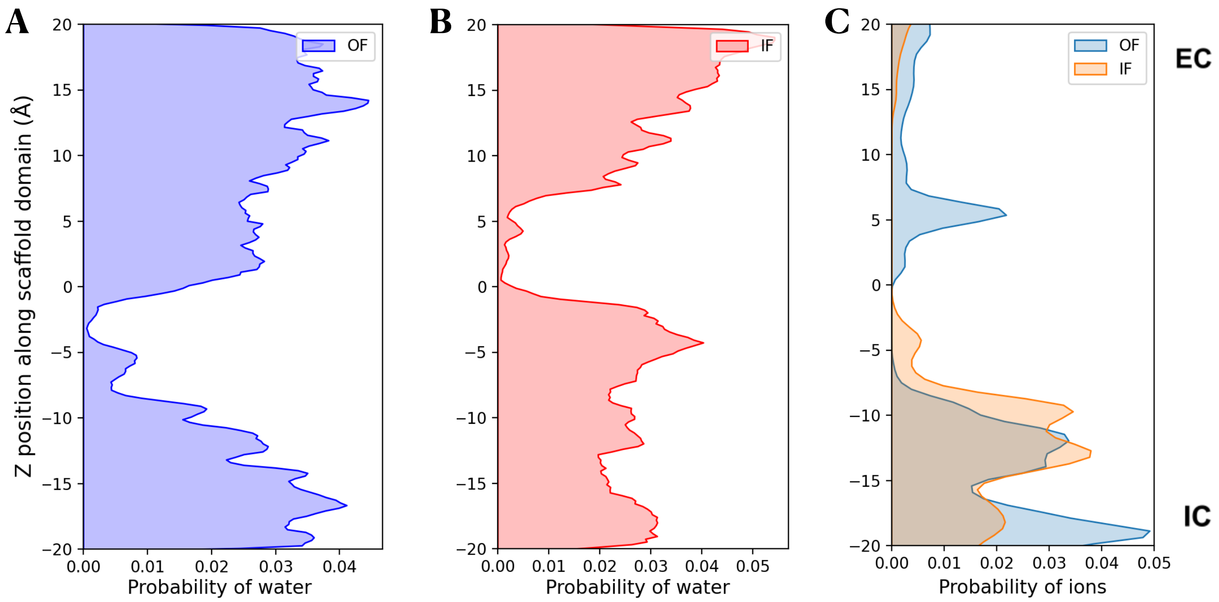


Figure S3: **Water and ion density in the AE1 lumen.** (A) Hydration probability of the lumen along the $z$-axis in the OF state during the equilibrium simulation. (B) Hydration probability in the IF state. (C) Anion distribution along the $z$-axis in both OF and IF states aligned using the SD, with the negative side representing the cytoplasm.

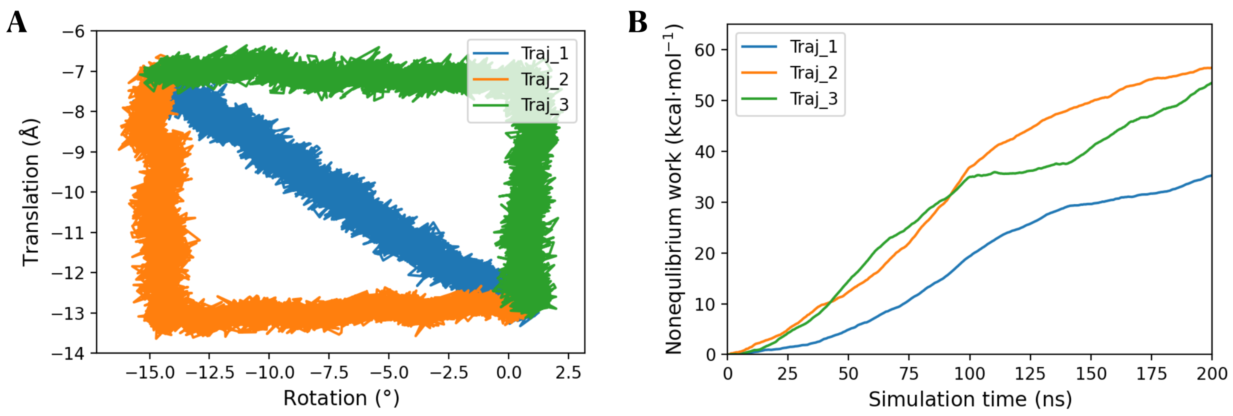


Figure S4: **Different pulling protocols and associated non-equilibrium work.** (A) Three different pulling protocols were tested for transition from OF to IF, using the same simulation times and force constants: (1) first rotating and then translating (green), (2) pulling in the reverse order (orange), and, (3) pulling synchronously (blue). (B) The non-equilibrium work generated from the three pulling protocols.

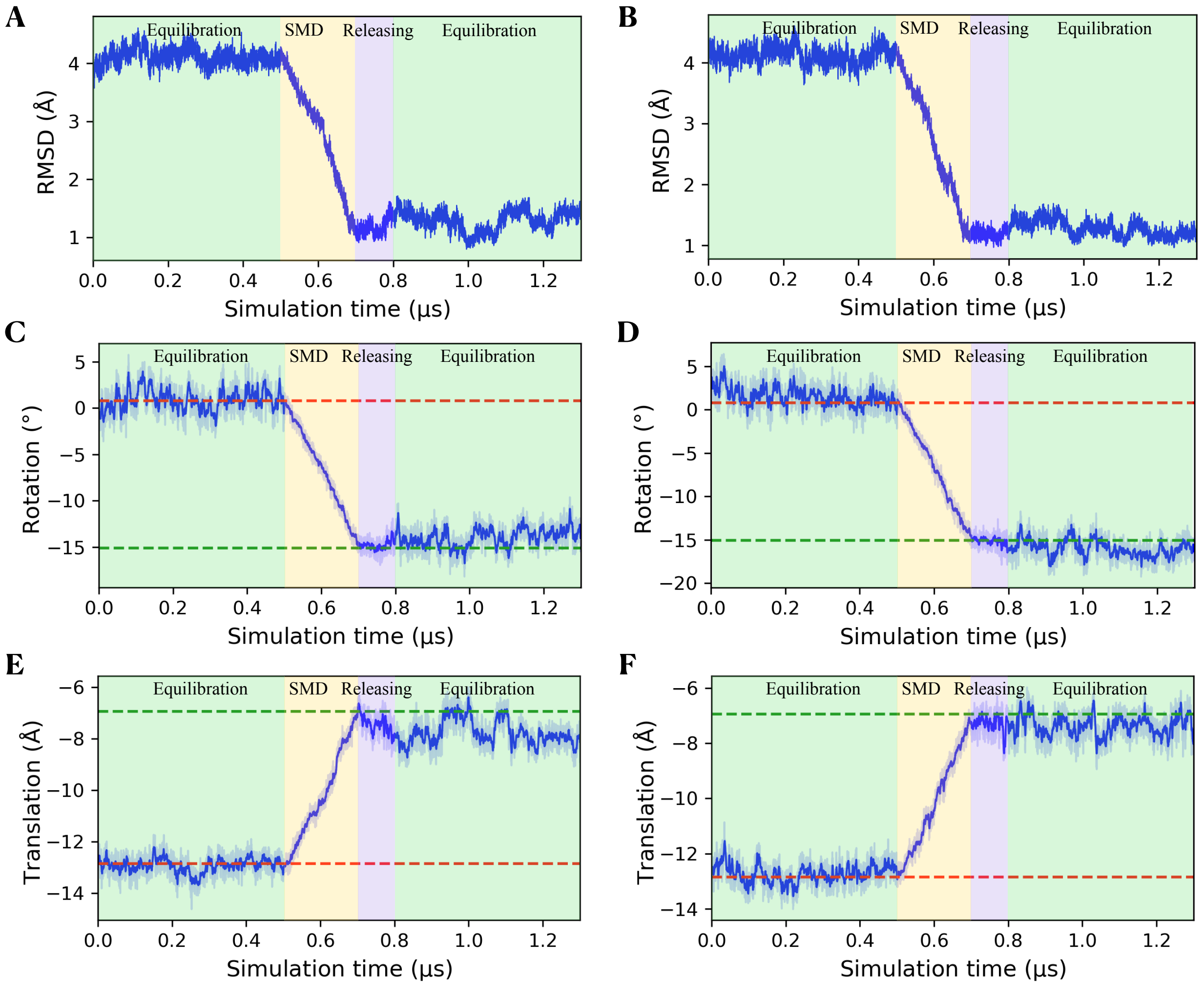


Figure S5: **RMSD and CV changes during the transition simulations of the *apo* and the substrate-bound PIP$_2$-removed AE1 systems.** (A and B) C$\alpha$ RMSD of TM helices of the TD during the driven simulation with respect to the target structure in the IF state of the *apo* (A) and the substrate-bound, PIP$_2$-removed (B) systems. (C and D) The TD rotation CV changes during the equilibration, driven, releasing, and final free equilibration stages for the *apo* (C) and the PIP$_2$-removed (D) systems. (E and F) The TD translation CV changes of the *apo* (E) and the PIP$_2$-removed (F) systems.

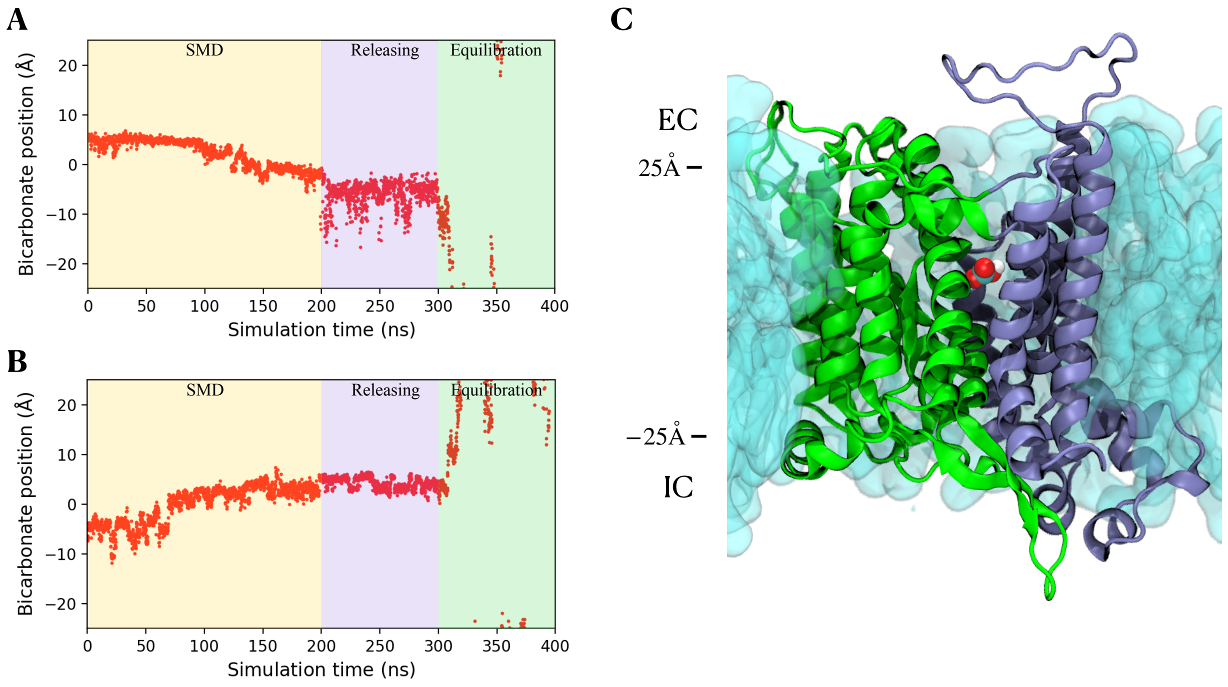


Figure S6: **$HCO_3^-$ transport in AE1 during state transition.** (A) $z$ coordinate of $HCO_3^-$ during the state transition from OF to IF2, releasing, and final free equilibration stages. The frames are aligned using the SD. (B) $z$ coordinate of $HCO_3^-$ during the reverse process (IF2 to OF transition). (C) Transmembrane spanning of AE1 (numbers matching those in Panel B. The membrane spans from −25 Å to 25 Å. The bound $HCO_3^-$ is located at $z$ = 5 Å in the OF state.

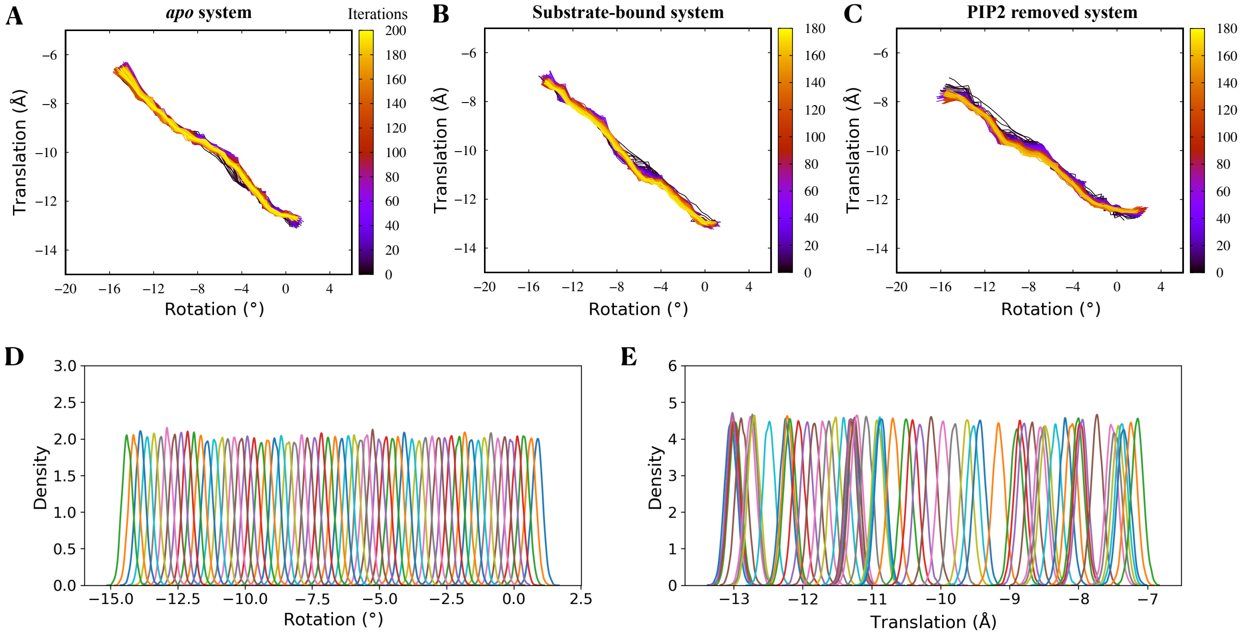


Figure S7: **Convergence of enhanced sampling simulations.** (A) The refinement of the transition pathway during 200 iterations of SMwST simulations for the *apo* system. The color is changing from black to yellow to show the iterations. (B) The pathway refinement for the substrate-bound system. (C) The pathway refinement for the substrate-bound, $PIP_2$-removed system. (D) The histograms for the rotation CV from BEUS simulations. The density curve of each window is calculated using the Guassian KDE function. (E) The histograms for the translation CV.

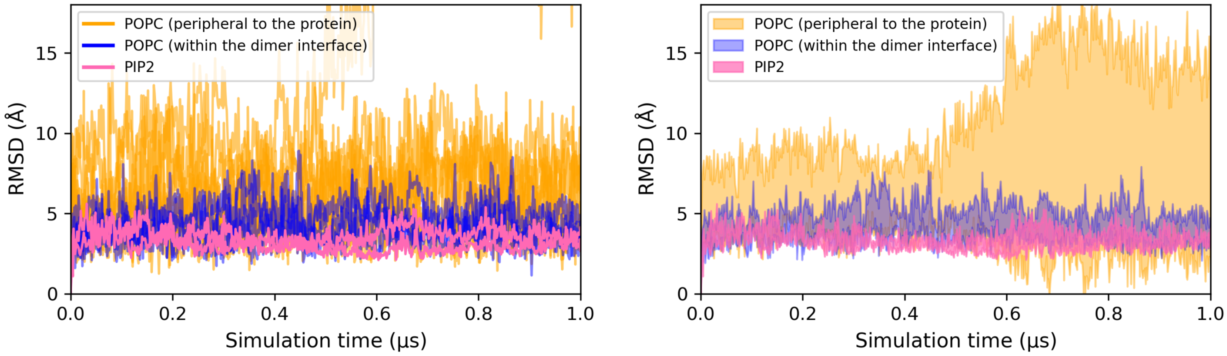


Figure S8: **RMSD of cryo-EM resolved lipids in MD.** (A) Headgroup RMSD during the 1 µs equilibrium simulation for the three groups of lipids relative to their initial positions are shown: 8 peripheral POPC lipids (orange), 4 POPC lipids within the dimer interface (blue), and 2 $PIP_2$ lipids (pink). (B) Standard deviation of the RMSD values for the same group, indicating the variability over the simulation time.

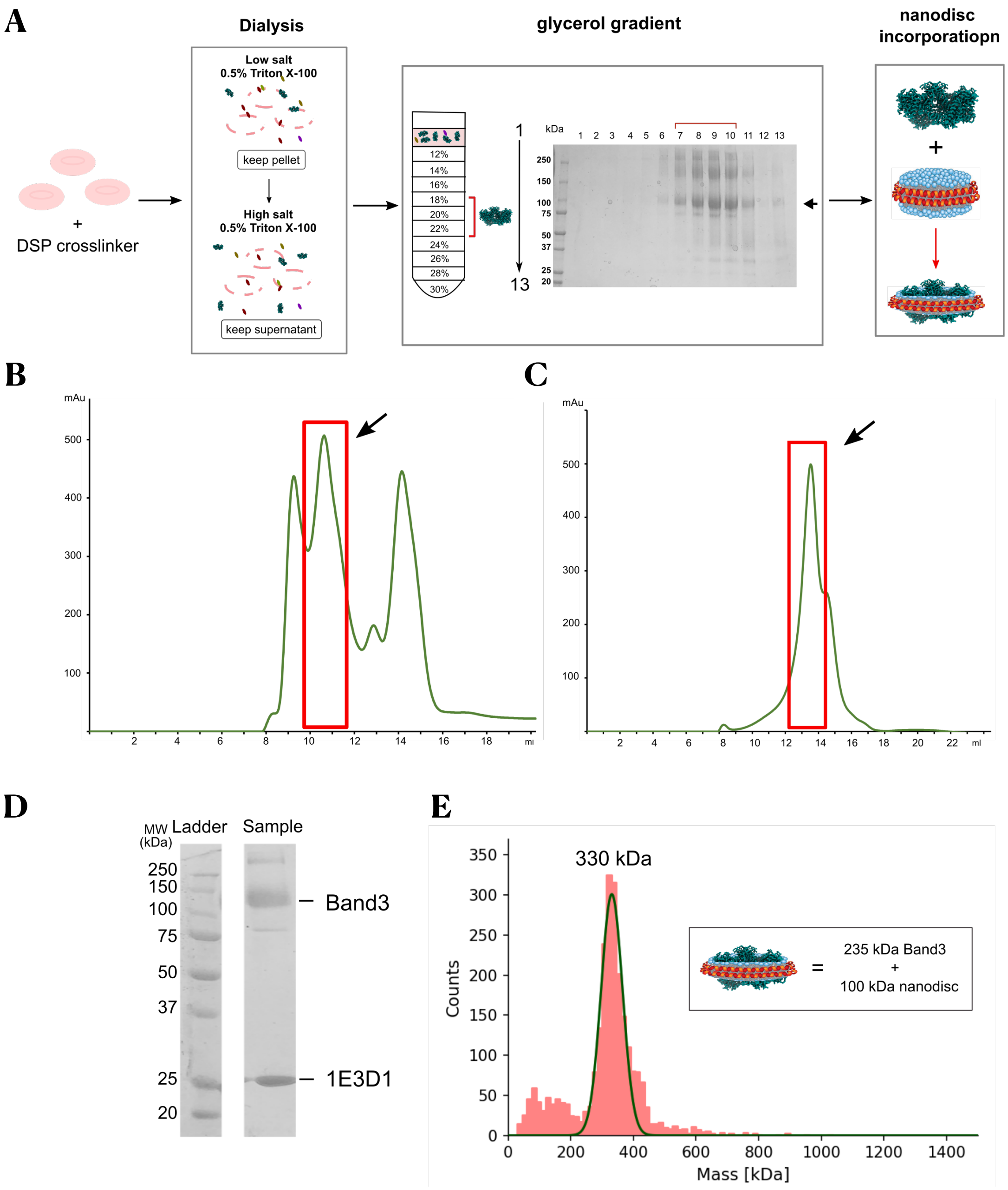


Figure S9: **Purification and biochemical characterization of human AE1.** (A) Human erythrocyte ghost membranes, represented as pink empty erythrocytes, were washed and solubilized using 0.5% (w/v) Triton-X100. Membrane fragments were then added in a 12-30% glycerol gradient. Fractions collected were loaded in an SDS-PAGE gel. Gradient fractions containing AE1 are marked with a red line. AE1 was incorporated in MSP nanodiscs. (B) Size exclusion chromatography of the fractions containing AE1 in nanodisc using a Superdex 200 column. The second peak (marked with the red box) was collected. (C) Collected fractions were subjected to a second round of size exclusion chromatography using a Superose 6 column. (D) SDS-PAGE gel of fraction that corresponds to the main peak from the gel filtration in panel C. (E) Mass photometry analysis of the size exclusion chromatography peak from the second gel filtration. The peak corresponds to a population of particles with a size around 330 kDa that represents the AE1 dimer in MSP nanodisc.

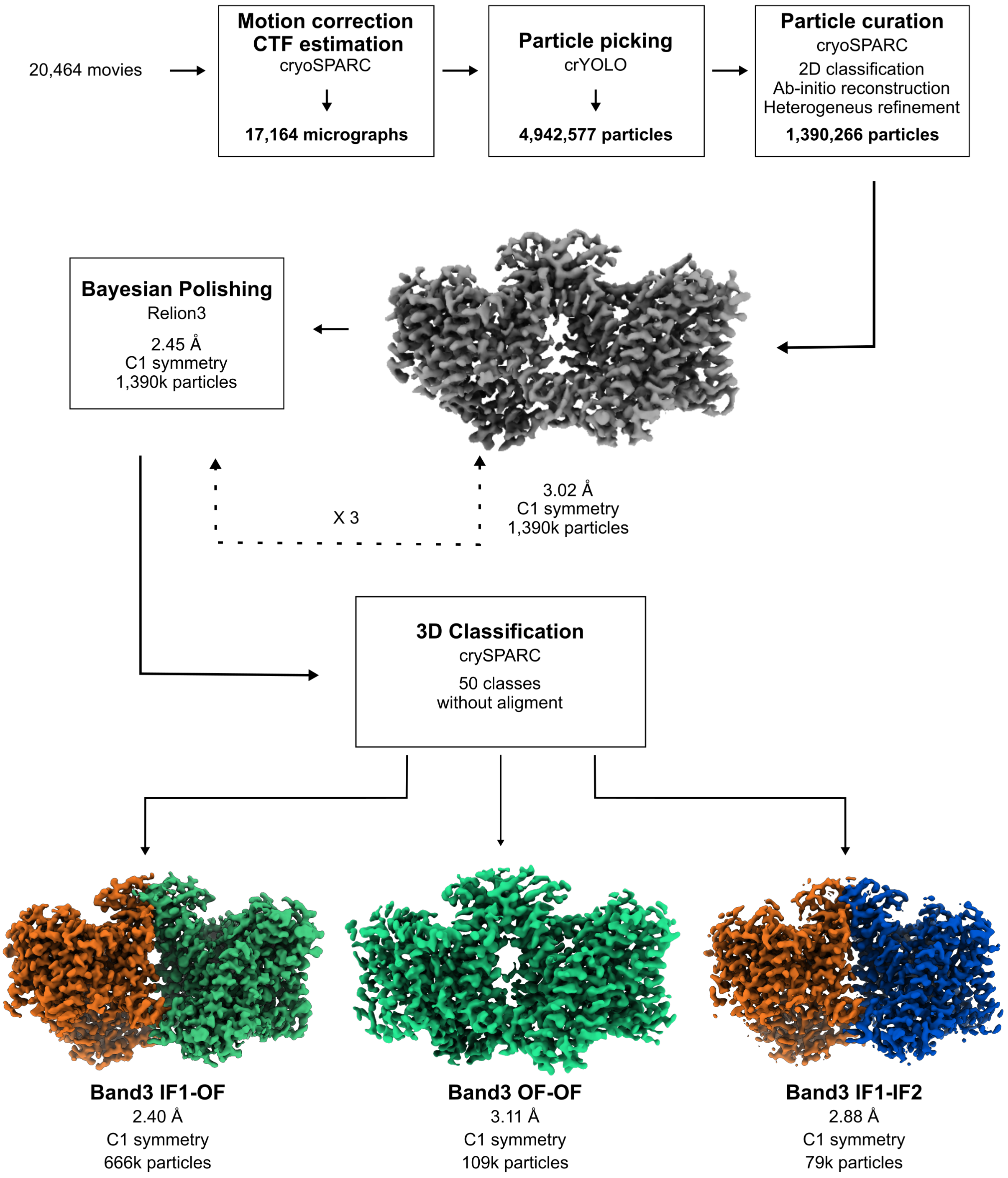


Figure S10: **Cryo-EM workflow.** Flowchart outlining cryo-EM image acquisition and processing performed to obtain the structure of AE1 in three different conformations. All processing was performed using CryoSPARC v.3.2 and Relion 4 (see Methods for details).

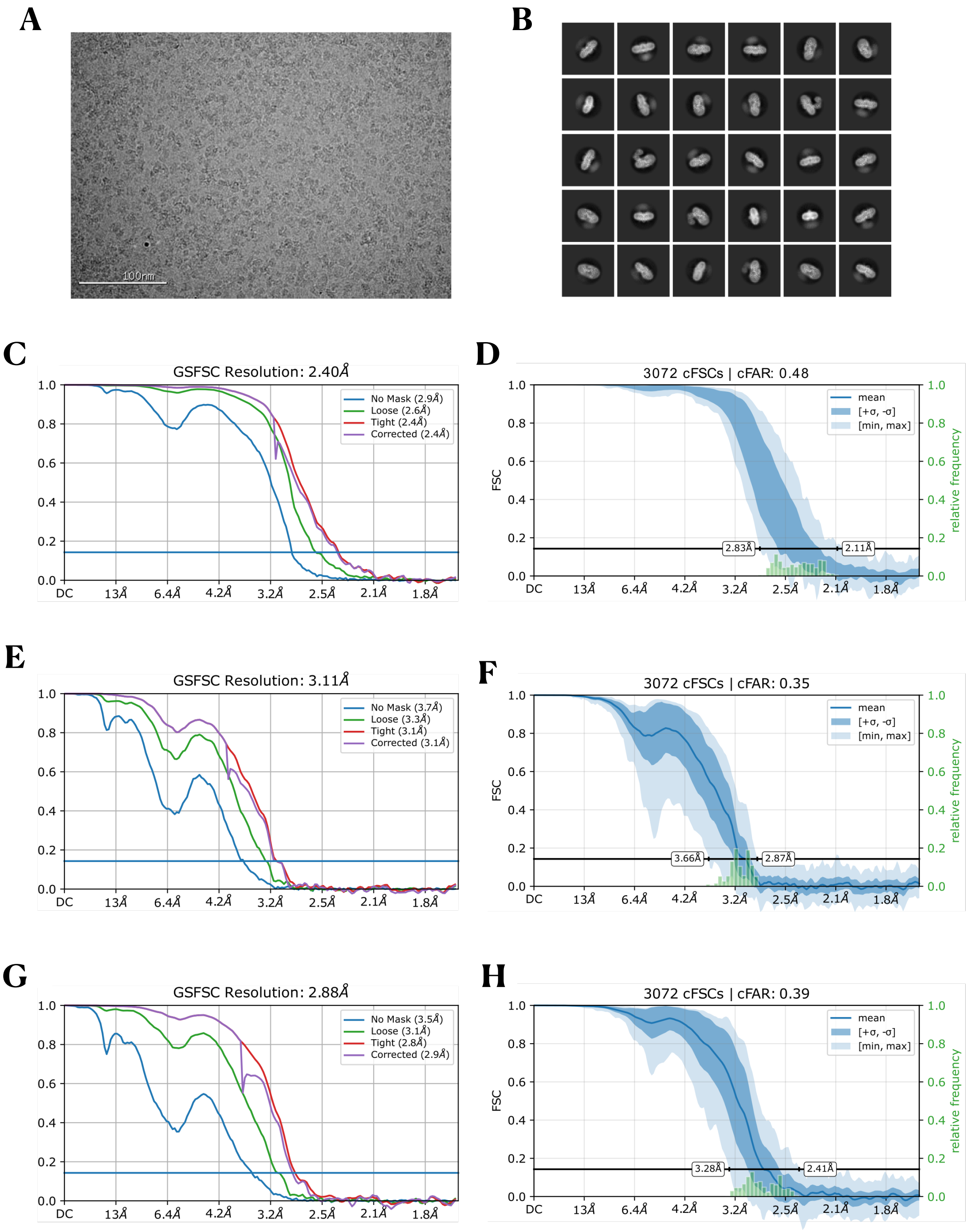


Figure S11: **Single particle cryo-EM structure determination.** (A) A representative micrograph. (B) 2D class averages of particles initially picked with crYOLO (~1.3M),[52] ordered by population. (C) Fourier shell correlation (FSC) curves for the AE1 dimer in the IF1-OF structure. (D) Directional anisotropy of the IF1-OF structure, as calculated by Orientation Diagnostics in CryoSPARC 4.5. (E) FSC curves for the AE1 dimer in OF-OF structure. (F) Directional anisotropy of the OF-OF structure, as calculated by Orientation Diagnostics in CryoSPARC 4.5. (G) FSC curves for the AE1 dimer in IF1-IF2 structure. (H) Directional anisotropy of the IF1-IF2 structure, as calculated by Orientation Diagnostics in CryoSPARC 4.5.

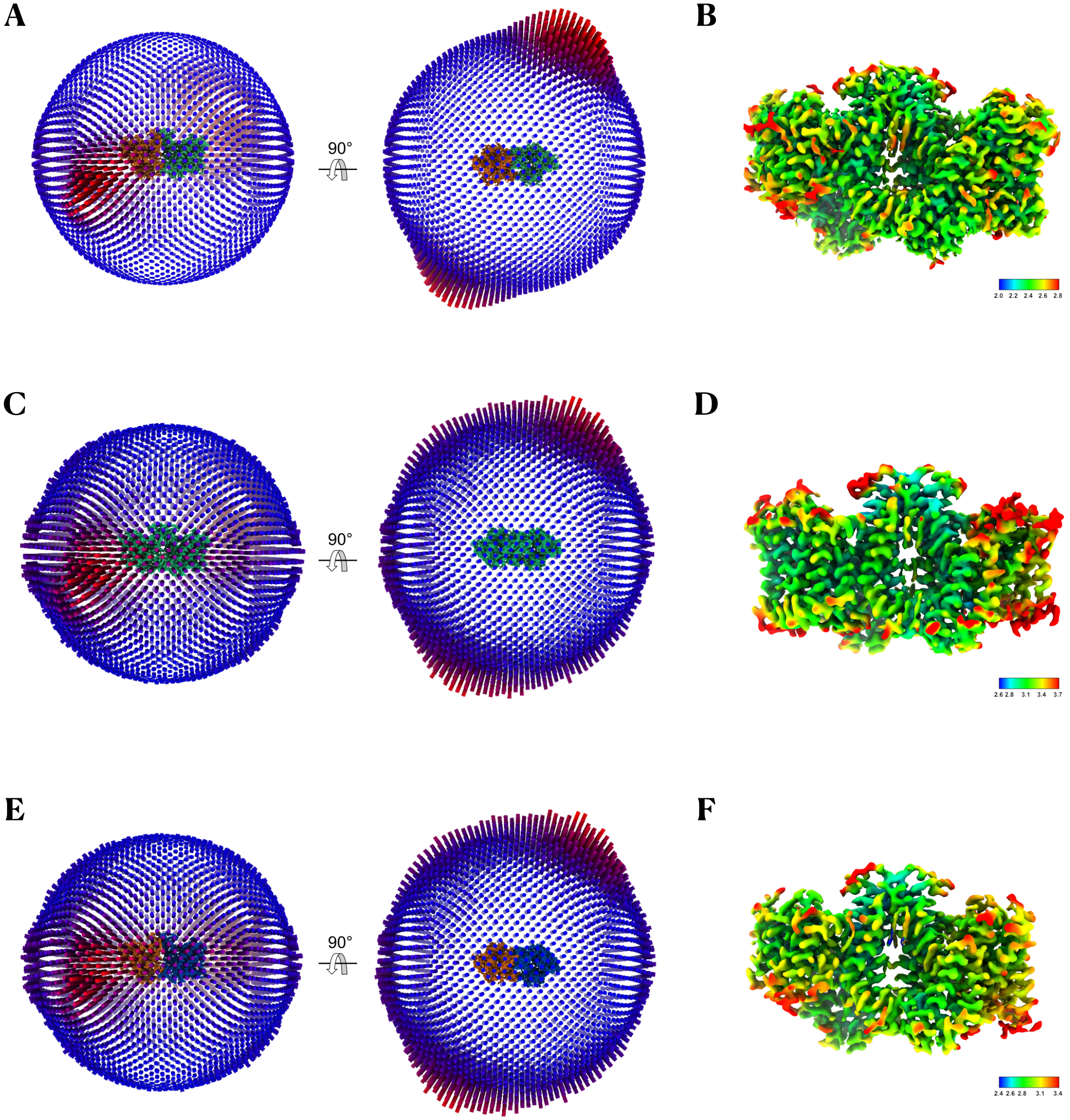


Figure S12: **Cryo-EM map plots.** (A) Orientation distribution plot for the 666,254 particles used in the IF1-OF refinement. (B) Local resolution plot of the cryo-EM map from the global 3D reconstruction for AE1 IF1-OF structure. (C) Orientation distribution plot for the 108,734 particles used in the OF-OF refinement. (D) Local resolution plot of the cryo-EM map from the global 3D reconstruction for AE1 OF-OF structure. (E) Orientation distribution plot for the 78,596 particles used in the AE1 IF1-IF2 refinement. (F) Local resolution plot of the cryo-EM map from the global 3D reconstruction for AE1 IF1-IF2 structure.

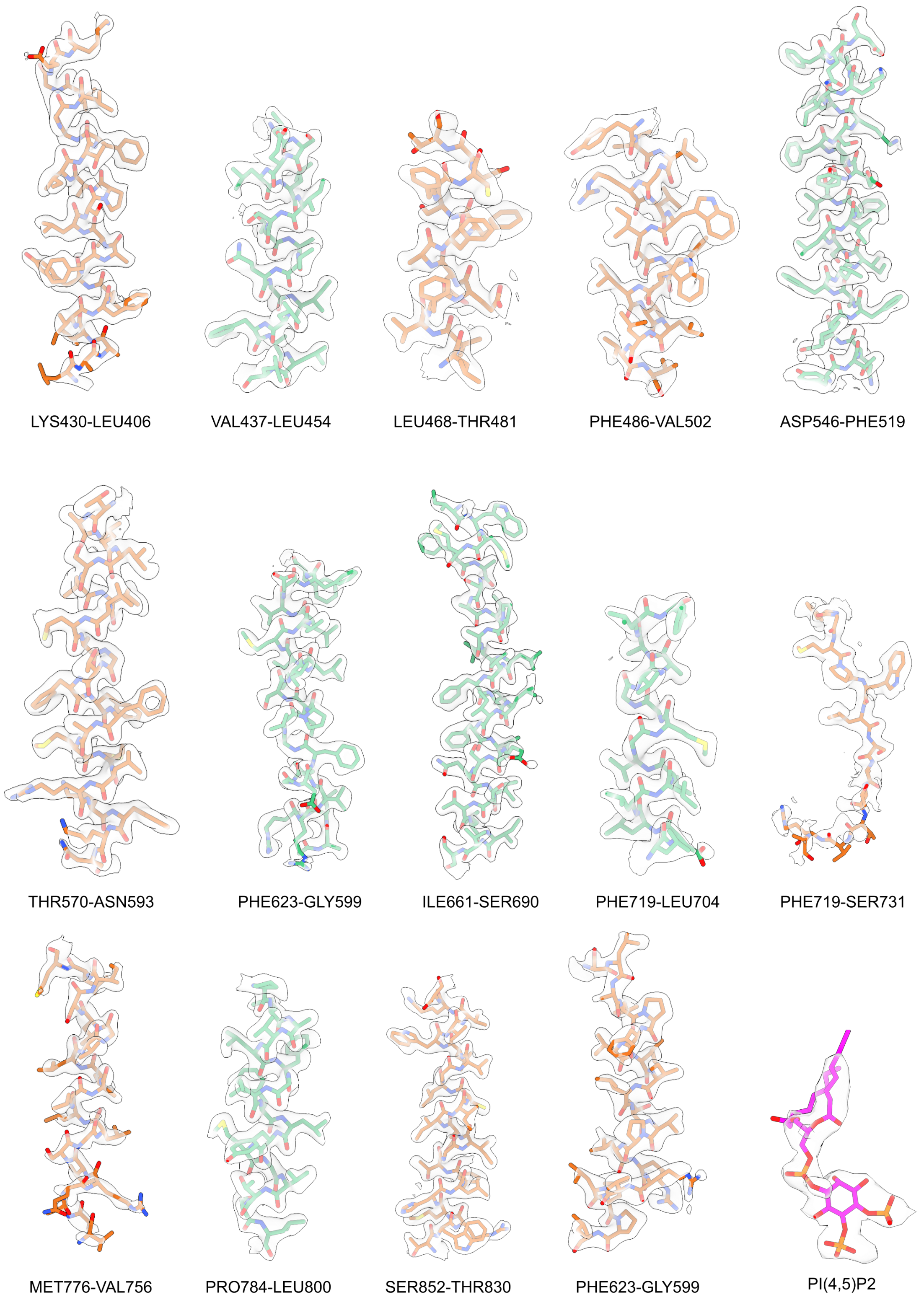


Figure S13: **Model/map fit.** Cryo-EM densities (transparent gray surface) are shown with corresponding segments of the AE1 IF1-OF atomic model; side chains are rendered in stick representation and colored as in Fig. 1.

#### Supplementary Table

| | AE1 $IF_1$-OF<br>PDB 9MND<br>EMD-48421 | AE1 OF-OF<br>PDB 9MNG<br>EMD-48422 | AE1 $IF_1$-$IF_2$<br>PDB 9MOS<br>EMD-48480 |
|---|---|---|---|
| **Data collection and processing** | | | |
| Magnification | 105,000x | 105,000x | 105,000x |
| Voltage (kV) | 300 | 300 | 300 |
| Electron exposure ($e^-$/Å$^2$) | 58 | 58 | 58 |
| Defocus range (µm) | -0.5 to -1.5 | -0.5 to -1.5 | -0.5 to -1.5 |
| Pixel size (Å) | 0.83 | 0.83 | 0.83 |
| Symmetry imposed | C1 | C1 | C1 |
| Initial micrographs | 20,087 | 20,087 | 20,087 |
| Final micrographs | 17,164 | 17,164 | 17,164 |
| Initial particle images | 1,781,329 | 1,781,329 | 1,781,329 |
| Final particle images | 666,254 | 108,734 | 78,596 |
| Map resolution (Å) | 2.4 | 3.11 | 2.88 |
| FSC threshold | 0.143 | 0.143 | 0.143 |
| **Model composition** | | | |
| Non-hydrogen atoms | 8519 | 8310 | 8240 |
| Protein residues | 982 | 1034 | 971 |
| Ligand molecules | 26 | 8 | 18 |
| **RMS Deviations** | | | |
| Bond lengths (Å) | 0.008 | 0.002 | 0.005 |
| Bond angles (º) | 1.222 | 0.493 | 0.885 |
| **Validation** | | | |
| Clashscore | 2.77 | 4.5 | 5.53 |
| Poor rotamers (%) | 2.42 | 2.93 | 0.97 |
| **Ramachandran plot** | | | |
| Favored (%) | 97.22 | 98.34 | 97.18 |
| Allowed (%) | 2.57 | 1.81 | 2.82 |
| Disallowed (%) | 0.21 | 0 | 0 |
| **EMRinger Score** | 3.89 | 4.44 | 3.55 |

Table S1: **Cryo-EM data collection, refinement and validation statistics**
